# The *Daphnia* carapace and the origin of novel structures

**DOI:** 10.1101/2021.09.29.462403

**Authors:** Heather S. Bruce, Nipam H. Patel

## Abstract

Understanding how novel structures arise is a central question in evolution. Novel structures are often defined as structures that are not derived from (homologous to) any structure in the ancestor^1^. The carapace of the water flea *Daphnia magna* is a bivalved “cape” of exoskeleton that has been proposed to be one of many novel arthropod structures that arose through repeated co-option of genes that also pattern insect wings^2–4^. To determine whether the *Daphnia* carapace is a novel structure, we compare the expression of *pannier, araucan*, and *vestigial* between *Daphnia, Parhyale*, and *Tribolium*. Our results suggest that the *Daphnia* carapace did not arise by co-option, but instead derives from an exite (lateral lobe) that emerges from an ancestral proximal leg segment that was incorporated into the *Daphnia* body wall. The *Daphnia* carapace therefore appears to be homologous to the *Parhyale* tergal plate and the insect wing^5^. Remarkably, the *vestigial*-positive region that gives rise to the *Daphnia* carapace appears to be present in *Parhyale*^6^ and *Tribolium* as a small, inconspicuous protrusion. Similarly, the *vestigial*-positive regions that form thoracic tergal plates in *Parhyale* appear to be present in *Daphnia*, even though *Daphnia* does not form thoracic tergal plates. Thus, rather than a novel structure resulting from gene co-option, the *Daphnia* carapace appears to have arisen from a shared, ancestral tissue (morphogenetic field) that persists in a cryptic state in other arthropod lineages. Cryptic persistence of unrecognized serial homologs may thus be a general solution for the origin of novel structures.

## Results

Many crustaceans, such as crayfish, crabs, ostracods, and tadpole shrimp, have a carapace. Classical morphological studies have argued that the carapace emerges from the dorsal posterior of the head (at approximately maxilla 2)^7,8^. In the water flea *Daphnia magna*, the carapace is a bivalved “cape” of exoskeleton that surrounds the animal. Shiga 2017 ^2^ confirmed the classical idea that the *Daphnia* carapace emerges from the dorsal posterior region of the head and, like insect wings, is composed of a bilayered sheet of ectodermal cells and expresses and requires the “wing” genes *wingless* (*wg*), *vestigial* (*vg*) and *scalloped* (*sd*)^2^. They propose that the *Daphnia* carapace and other flat, lateral lobes in arthropods arose by multiple instances of co-option of wing patterning genes.

However, the analyses in Bruce and Patel 2020^5^ and Bruce and Patel 2021^9^ suggests an alternative hypothesis. These analyses, drawing on over a century of morphological and embryological studies as well as gene expression and loss-of-function studies, generated a model for understanding the homologies of any structure on any arthropod leg (Fig. 1). In this model, most arthropods have incorporated one or two ancestral proximal leg segments into the body wall ^5,9–13^ (leg segments 7 and 8, counting from the terminal claw), but the division between “true” body wall (tergum) and the incorporated leg segments that now function as body wall can be distinguished by the expression of *pannier* (*pnr*) and *Iroquois* complex genes such as *araucan* (*ara*). In the embryos of *Drosophila* (fruit fly), *Tribolium* (flour beetle) *Parhyale* (amphipod crustacean), and *Acanthoscurria* (tarantula), *ara* expression brackets the hypothesized incorporated 8^th^ leg segment, while *pnr* is expressed in the dorsal-most tissue and marks the true body wall^9^. Bruce and Patel 2020^5^ showed that the insect wing and *Parhyale* tergal plate are both derived from ancestral exites: multi-functional lobes that emerge from proximal leg segments and that express and require “wing” genes such as *vg* and *sd*^6,14,15^. They showed that the insect wing and *Parhyale* tergal plate emerge from the ancestral leg segment 8 that was incorporated into the body wall (Fig. 1B).

**Fig.1.**
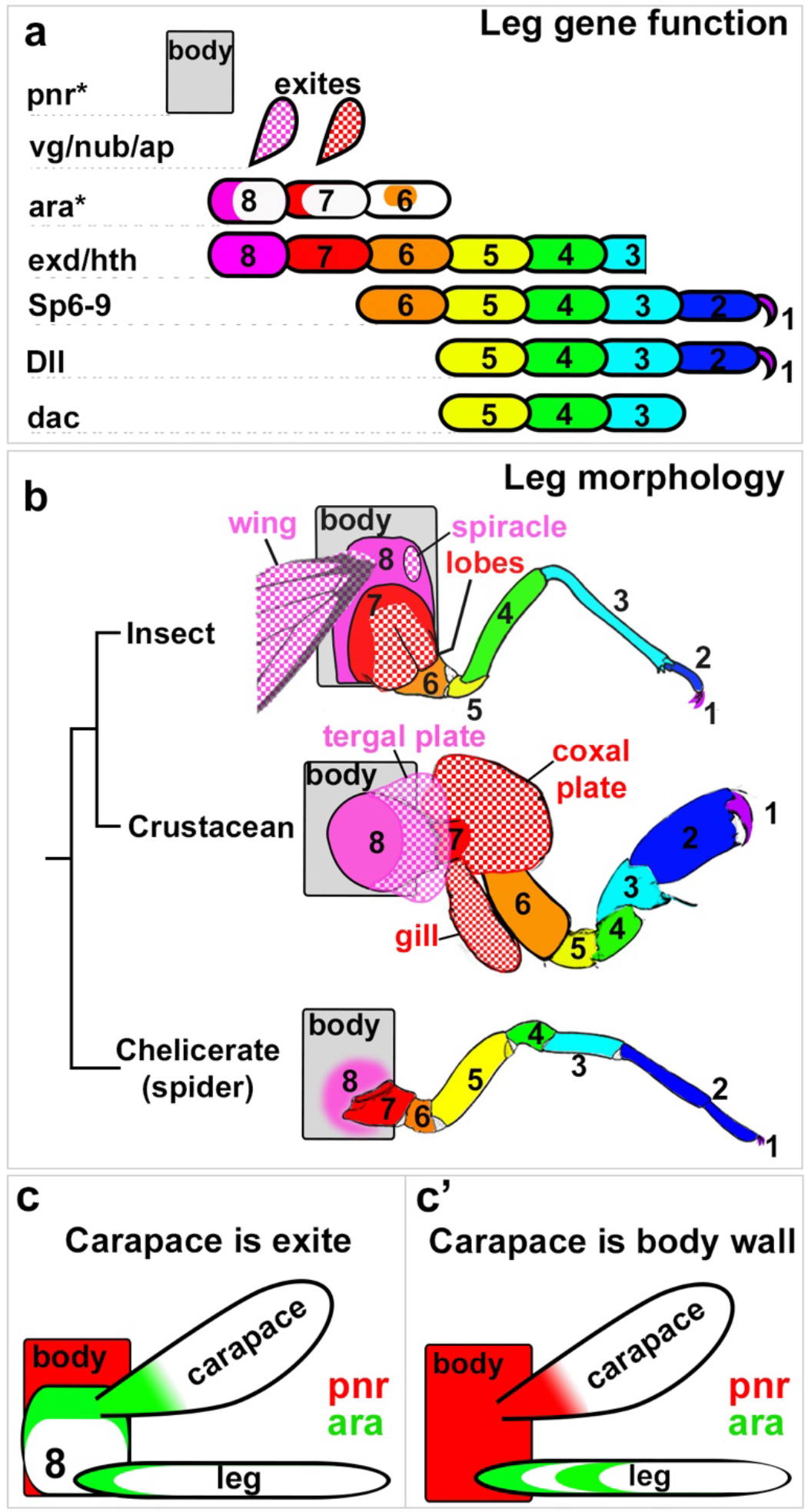
Model of how to align all arthropod legs from Bruce 2021 ^9^. **a**. A schematic of the leg structures that require each gene in chelicerates, crustaceans, insects provides a model for how to align crustacean and insect legs. Based on the function of *exd, hth, Dll, Sp6-9*, and *dac*, the six distal leg segments (leg segment 1 through leg segment 6) of chelicerates, crustaceans, and insects correspond with each other in a one-to-one fashion. The alignment of the two proximal leg segments is based on expression of *pnr* and *ara* in chelicerates, crustaceans, and insects, and the function of wing/exite genes in insects and crustaceans. **b**. Morphology and proposed homologies of arthropod leg segments. Colors and patterns indicate proposed homologies. Exites (checker pattern); endites (stripe pattern). Drawings in b modified from Snodgrass 1952. Reproduced from Bruce & Patel 2021. **c-c’**. Based on the above model, predictions can be made about the expression of *pnr* (red) and *ara* (green) in the *Daphnia* carapace. If the carapace is an exite on an incorporated 8^th^ leg segment (c), *pnr* will be expressed in a narrow stripe dorsal to the carapace and the *ara* domain adjacent to *pnr* will extend into the carapace. If the carapace is an outgrowth of the body wall (c’), then *pnr* will extend into the carapace, and the two domains of *ara* will be located ventral to the carapace.

Based on this model, the morphological and molecular data in Shiga 2017 suggests that the *Daphnia* carapace did not arise by co-option, but instead represents an exite on an incorporated 8^th^ leg segment of the head. The *Daphnia* carapace would therefore be homologous to the *Parhyale* tergal plate^5,6^ and the insect wing^5^.

To test the proximal-distal register of the *Daphnia* carapace, the expression of *pannier*, an *Iroquois* gene, and the wing/exite patterning gene *vg* was examined in embryos of *Daphnia magna, Tribolium castaneum*, and *Parhyale hawaiensis* using in situ hybridization chain reaction (HCR) version 3.0^16,17^. A single *pnr* gene was identified in *Daphnia* which was the reciprocal best blast hit of *Drosophila, Tribolium*, and *Parhyale pnr* ^5^. A single *Iroquois* complex gene was identified in *Daphnia* which was the reciprocal best blast hit of *Drosophila, Tribolium*, and *Parhyale ara*. This *Daphnia* gene is hereafter referred to as *ara* ^5^. *Daphnia vg* was identified previously by Shiga 2017^2^.

If the *Daphnia* carapace is the exite of the incorporated 8^th^ leg segment, then our model predicts that *pnr* will be expressed in a narrow stripe dorsal to the carapace and the *ara* domain adjacent to *pnr* will extend into the carapace. Alternatively, if the carapace is a dorsal, non-leg-derived structure, then *pnr* expression should extend into the carapace, and the two domains of *ara* will be located ventral to the carapace. In either case, *vg* will be expressed along the edge of the carapace^2^.

Consistent with the hypothesis that the carapace is an exite on leg segment 8, *Daphnia vg* is expressed along the edge of the carapace, *pnr* is restricted to a narrow, dorsal stripe above the carapace, and the *ara* domain adjacent to *pnr* extends into the carapace (Figs. 2 and 3). Interestingly, *ara* expression on leg segment 6 appears to mark the position of the exopod. In a uniramous *Parhyale* leg (Figs. 3e) that lacks an exopod, the leg segment 6 *ara* domain forms a circular patch on the lateral side of the leg. However, in the biramous (split) legs of *Daphnia* (Figs. 2d and 3f, S1) and *Parhyale* (Fig. S2) where an exopod emerges on the lateral side of leg segment 6, the *ara* patch is expanded, and in *Parhyale*, the patch encircles the base of the exopod (see Fig. S1 for explanation of *Daphnia* leg identification).

**Fig.2.**
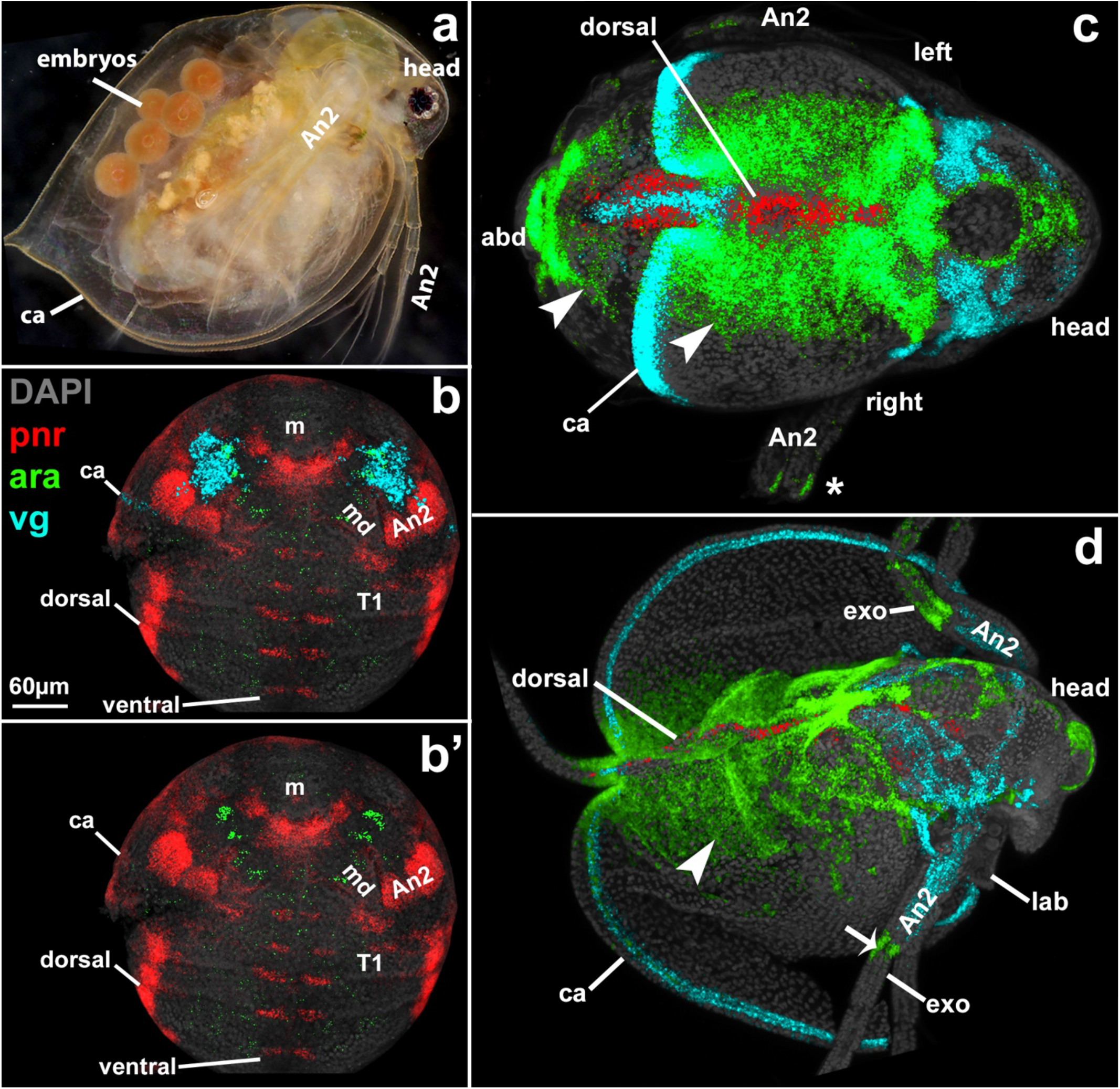
Expression of *pannier, araucan*, and *vestigial* in *Daphnia* embryos. **a**. *Daphnia* adult with embryos under carapace (ca), lateral view. **b, b’**. Stage 7 embryo, ventral view. Maxilla 1 and 2 not yet visible but will emerge between mandible (md) and first thoracic leg (T1) **c**. Stage 9 embryo, dorsal view. **d**. Dissected head and carapace of St 11 embryo, dorsal view, showing *ara* and *pnr* expression without trunk underneath. *pnr* marks the dorsal-most domain. *ara* is expressed in four domains: in a dorsal region adjacent to *pnr* (closed arrowhead), in a second domain on leg segment 7 (not visible here), at the base (arrow) of the exopod (exo, on leg segment 6 in uniramous legs), and in the tip of leg segment 1 (only present in antenna 2 (An2). *vg* is expressed in the perimeter of the carapace (ca), and in the mesoderm of antenna. m, mouth. abd, abdomen. *pnr* (red), *ara* (green), *vg* (light blue), DAPI (grey).

**Fig.3.**
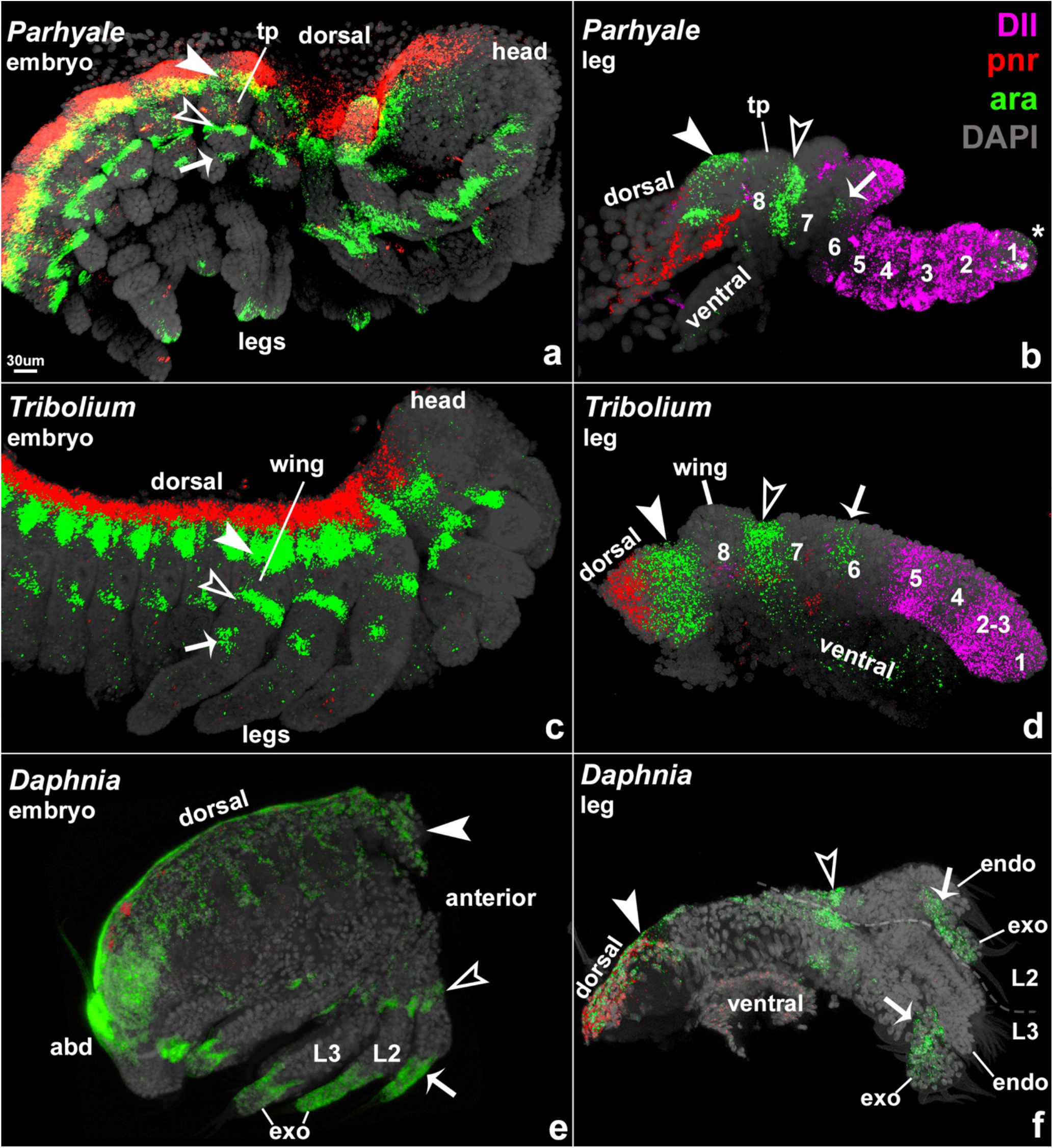
Expression of *pnr* and *ara* elucidates the proximal leg segments. Dissected right half of *Parhyale* (a), *Tribolium* (c), and *Daphnia* (e) embryos. *Daphnia* head and carapace removed to reveal trunk. Dissected legs of *Parhyale* (b), *Tribolium* (d), and *Daphnia* (f) embryos. Large cells dorsal to *pnr* expression in a and b are extra-embryonic cells that exist prior to dorsal closure. *pnr* marks the dorsal-most domain. *ara* is expressed in four domains: a dorsal region adjacent to *pnr* (closed arrowhead); a second region on leg segment 7 (open arrowhead); a third region on leg segment 6 (arrow); and, in crustaceans and tarantula^9^, in the tip (*) of leg segment 1. Leg segment 6 domain is a dot when no exopod (exo) is present (arrow in e). Dot becomes expanded in exopod when leg is biramous (arrows in d). Note that *ara* expression in e and f in exopod does not extend to tip as it does in a and b, therefore this exopod domain is not the leg segment 1 (^*^) domain. Indeed, *Daphnia* thoracic legs are truncated and do not have leg segment 1, see Fig. S1 for explanation of *Daphnia* leg identification. However, *Daphnia* antenna 2 has this domain (Fig. 2d). *pnr* and *ara* are expressed in a smattering of ventral non-leg cells. tp, tergal plate. b, c, e, f from Bruce 2020 and Patel^5^.

If the *Daphnia* carapace is the exite of the incorporated 8^th^ leg segment of a mouthpart (modified leg) on the head, this exite may still exist on the head appendages of arthropods that do not form a carapace. In support of this hypothesis, *vg* is expressed in the head of *Tribolium* dorsal/proximal to the mouthparts (Fig. S3A). This *vg* domain is bracketed by *ara* expression, just like the insect wing, the *Parhyale* tergal plate, and the *Daphnia* carapace. This region is therefore presumably homologous to the 8^th^ leg segment. Notably, there is no obvious structure associated with the mouthpart *vg* domain. In *Parhyale, vg* patterns the flange-like protrusion that protects the mouthparts, because the flange is reduced when *vg* is knocked out (Fig. S3B, C^6^). This flange emerges from the incorporated 8^th^ leg segment, because it is bracketed by *ara* expression. Given that the arthropod head is composed of several modified legs fused into a head capsule^18^, the head flange likely represents several adjacent exites. Thus, rather than new, co-opted domains of *vg* expression, these *vg* head domains are ancient and conserved.

Similarly, while *Daphnia* does not form thoracic tergal plates like *Parhyale*, the regulatory architecture for generating these structures still appears to exist in *Daphnia*. Shiga 2017 noted segmentally repeated dorsal domains of *wg* expression. These *wg* domains are co-expressed with *vg*. Like *Parhyale* tergal plates, this *Daphnia wg-vg* domain is ventral to *pnr* and bracketed by the two armbands of *ara*. Together, this molecular triangulation suggests that these segmentally repeating *wg-vg* domains in *Daphnia* are morphogenetic fields (a discrete set of cells capable of forming a specific organ, e.g., leg, wing, bristle, etc) that pattern exites such as thoracic tergal plates, just like the segmentally repeating *vg*-positive *Parhyale* tergal plates. Thus, even though *Daphnia* does not form tergal plates on the thorax, it still retains the regulatory architecture for making them in serially homologous positions in each thoracic segment. These thoracic exite fields could presumably be activated to form an obvious structure in future lineages.

Shiga et al. 2017^2^ note that the expression domains of *hedgehog* (*hh*)/*engrailed* (*en*) and *wg* in *Drosophila* wings and the *Daphnia* carapace are not oriented as expected if the two are homologous: in *Drosophila* wings, *hh/en* and *wg* are expressed orthogonally to each other, while in the *Daphnia* carapace, *hh/en* and *wg/vg* are expressed parallel to each other (discussed in Tomoyasu 2021^19^). Comparisons with *Parhyale* are informative in explaining these observations. In *Parhyale* tergal plates, here proposed to be positionally homologous to insect wings and the *Daphnia* carapace, *vg* is expressed along the posterior edge in a J-shape^6^. Thus, in *Parhyale, vg* expression in tergal plates runs both orthogonally and parallel to *en* expression (Fig. S4). If the posterior edge of a *Parhyale* tergal plate expanded posteriorly, then *en* and *wg* would be expressed in parallel in this expanded structure, as they are in the *Daphnia* carapace. On the other hand, if the ventral edge of a *Parhyale* tergal plate were to expand ventrally, then *en* and *wg* would be expressed orthogonally to each other, as they are in insect wings. Thus, the difference in the expression axes of *wg* and *en* between *Drosophila* and *Daphnia* may be a function of the direction in which the exite expanded, rather than a lack of homology. Intriguingly, the posterior spine of the *Daphnia* carapace, which expresses both *pnr* and *vg*, may be positionally homologous to the insect scutellum, which is also a dorsal posterior protrusion of the tergum that expresses *vg*^19–21^.

## Discussion

Taken together, the expression and knockdown data in Shiga 2017 and the expression data presented here suggest that the *Daphnia* carapace evolved by posterior expansion of the exite of the incorporated 8^th^ leg segment of maxilla 2. The *Daphnia* carapace would therefore be homologous to the *Parhyale* tergal plate and the insect wing (Fig. 4). If the *Daphnia* carapace is an exite, then the carapaces of crustaceans in general, which also appear to emerge from the dorsal-posterior head at mx2^7,8,22^, may also be exites.

**Fig.4.**
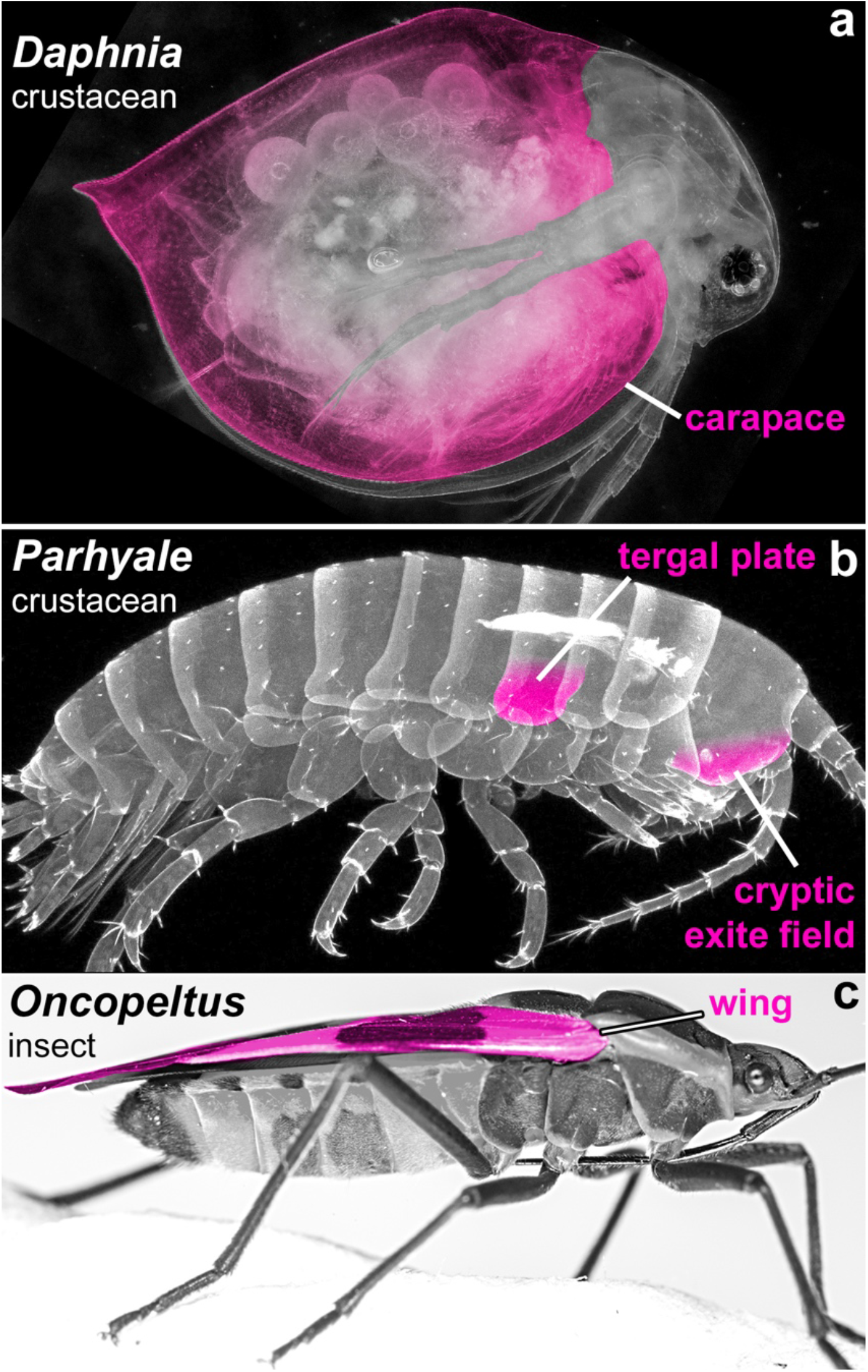
The *Daphnia* carapace is hypothesized to be the exite of the incorporated 8^th^ leg segment, homologous to the *Parhyale* tergal plate and insect wing. Homologous structures indicated with pink shading. Not all serially homologous structures in all animals are indicated.

Many arthropods have dorsal and lateral structures (carapaces, plates, gills, glands, horns, helmets, etc), the homologies of which are frequently debated^14,23^. The work here demonstrates that the proximal-distal identity of these structures can be elucidated using the expression of *pnr* and *Iroquois* genes such as *ara*, and further pinpointed with leg segment joint markers like *odd*-*skipped*, distal leg markers such as *Distal-less* (*Dll*), and exite genes such as *vg*. This simple molecular triangulation strategy, which does not require functional studies and can be accomplished using the affordable, forgiving, and straightforward in situ HCR v3.0^16,17^, can illuminate the homologies of long-debated structures on the legs and body wall of arthropods and determine which, if any, are truly novel.

Understanding how novel structures arise is a central question in evolution. Novelties are often defined as structures that are not homologous to any structure in the ancestor nor to any other structure in the same organism^1^. Arthropod legs and bodies are decorated with a fascinating diversity of structures, many of which have been proposed to be novel. Co-option of genetic pathways and “deep homology” has become a dominant explanation for the origin of novel structures within the field of evolutionary developmental biology (evo devo)^2–4,24–33^.

However, the work presented here, and in Bruce and Patel 2020^5^ and Bruce 2021^9^, provides an alternative hypothesis: these leg-associated structures (and perhaps other novel structures too) arise from unrecognized, serially homologous, morphogenetic fields (morphogenetic field = a discrete set of cells programmed to form a specific organ, e.g., leg, wing, bristle, etc). In this model, morphogenetic fields can persist in a cryptic, unrecognizable form (such as the *Parhyale* head flange, Figs. 4, S3) in intermediate lineages and become elaborated again in later lineages (such as the *Daphnia* carapace, Fig. 4), such that they may no longer be recognizable as the ancestral structure and appear to be novel. These conserved, cryptic fields are an expected outcome of the Character Identity Mechanisms (ChIMs) of DiFrisco et. al.^34^. Cryptic fields are similar to phyletic or taxic atavisms^35^, except that cryptic fields are not limited to reversions to ancestral structures, and may in fact still form a structure, but the structure is not recognized as homologous to the ancestral structure.

Fortunately, the complete palette of unrecognized serially homologous fields should be relatively straightforward to identify by employing experiments that transform homologous tissues into each other. For example, Hox knockout and misexpression studies in insects can cause ectopic legs or wings^36,37^. All of these ectopic structures are simply transformations of existing, serially homologous tissue along the anterior-posterior axis^36,38^. These homologous locations could be the source of future novel structures.

According to the cryptic persistence model, structures would not have to be continuously present in a morphologically obvious state from ancestor to descendant in order to be homologous. Indeed, a morphogenetic field may not form an obvious structure at all, as in the *vg*-positive regions of the *Tribolium* head (Fig. S3) and *Daphnia* dorsal body. Rather than de novo co-options, these morphogenetic fields are always there, persisting in a dormant, truncated, or highly modified state, and de-repressed in various lineages to form what look like novel structures. Cryptic persistence of morphogenetic fields may therefore provide a mechanistically satisfying explanation for the origin of novel structures.

## Acknowledgements

We thank Leon Peshkin for the *Daphnia* culture, Carsten Wolff for help with maintaining the culture. We thank Dennis A. Sun, Jennifer B. McCarthy, Sophia Kelly, Carsten Wolff, Yukimasa Kobayashi, Andrew Gillis, and Gregory Edgecomb for helpful comments. HSB thanks Lisa M Nagy for inspiring her quest for the origin of novel structures.

## Author contributions

HSB conceived, designed, and performed experiments and wrote manuscript. NHP edited manuscript.

## Star methods

### Animal care

*Daphnia* were kept in *Daphnia* culture medium^39^ in cleaned pickle jars with loose-fitting glass dishes for lids and fed daily with 3 – 6 drops of RGComplete (reefnutrition.com) depending on population size. To reduce overcrowding and resting egg production, all but the largest *Daphnia* were removed once every two weeks by pouring through two nets into a Tupperware, the first net with a 2mm pore size to catch the largest *Daphnia*, the second net with a fine pore size such that no hatchlings went through to the Tupperware. The 2mm net was then placed upside down over the mouth of the pickle jar and the water in the Tupperware was poured through the 2mm net, releasing the largest *Daphnia* back into the pickle jar.

### Fixation

From jars culled as above, the largest *Daphnia* were coaxed with light to one area of the pickle jar and removed with a plastic pipette with tip cut off to a Sylgard 184 (Dow Corning) dish.

Water was removed from Sylgard dish to immobilize Daphnia. *Daphnia* with embryos were picked up gently with forceps and placed in a medicine cup with culture medium. Once all animals with embryos had been gathered, animals were transferred to a 1.5mL Eppendorf tube, water removed with pipette, then animals were fixed for 1-2 hours by adding 3.2% aqueous paraformaldehyde (Electron Microscopy Sciences) in *Daphnia* culture medium. Less fixation time seems to reduce background. Fixed animals were washed 3×5min with PBS-Tween then dehydrated stepwise into methanol, then stored at -20C.

### In situ HCR

In situ HCR performed as in Bruce et al 2021^17^. Embryos were removed from adults during final PBS-Tween wash prior to pre-hybridization step.

### Imaging

Embryos imaged with Zeiss LSM880 confocal. Image processing done with Fiji-ImageJ. Fiji “Image Calculator > Subtract” method was used to remove high background from yolk autofluorescence. Figures processed using Adobe Photoshop 2020.

## Supplemental information legends

**Fig.S1.**
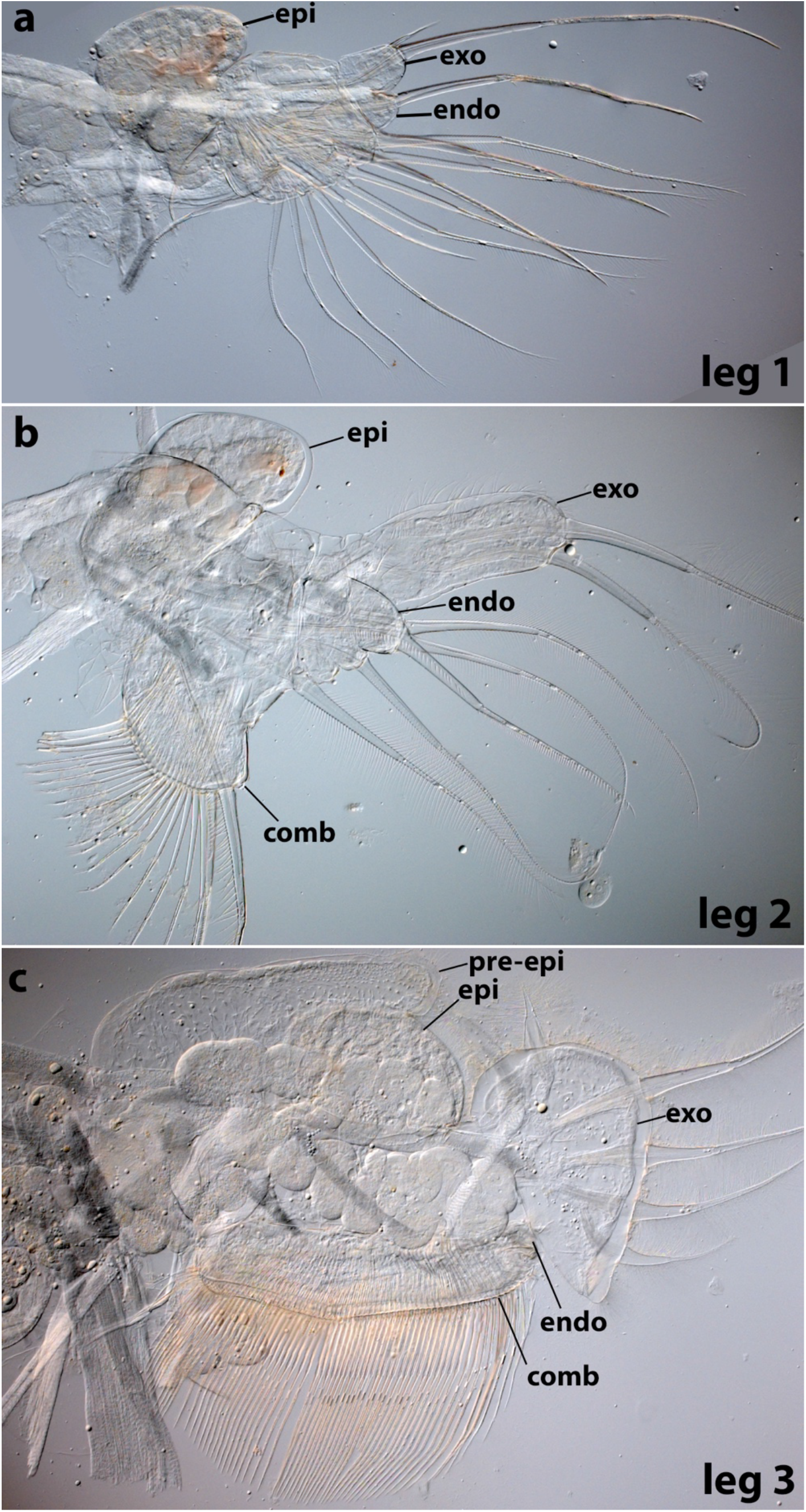
*Daphnia* adult dissected thoracic legs 1 - 3. *Daphnia* legs are “phyllopodous”, or leaf-shaped, and challenging to relate to walking legs in other arthropods such as *Parhyale* and *Tribolium*. However, a few key points facilitate comparisons. First, leg segment 6 can be identified because when crustacean legs are “biramous”, i.e., split into a lateral exopod (exo) and a medial endopod (endo), such as the *Daphnia* thoracic legs, then the exopod and endopod are carried on leg segment 6 (basis or basipod). Counting distally from leg segment 6, the endopod of leg 1 appears to have one segment and the endopod of leg 2 has at most 3 segments. Given that a complete endopod has 5 leg segments, and no terminal claw is apparent, *Daphnia* thoracic legs do not appear to express leg segment 1. Second, each *Daphnia* thoracic leg is identified by its shape and setal patterns. For example, Leg 2 has a short comb (gnathobase) and an elongated exopod with two long setae, while Leg 3 and 4 have a wide comb and a paddle-shaped exopod with an array of six setae. Endopod in c is underneath comb and not clearly visible. Epi, epipod (a type of exite) of the coxa (leg segment 7). Pre-epi, pre-epipod of leg segment 8^40^.

**Fig.S2.**
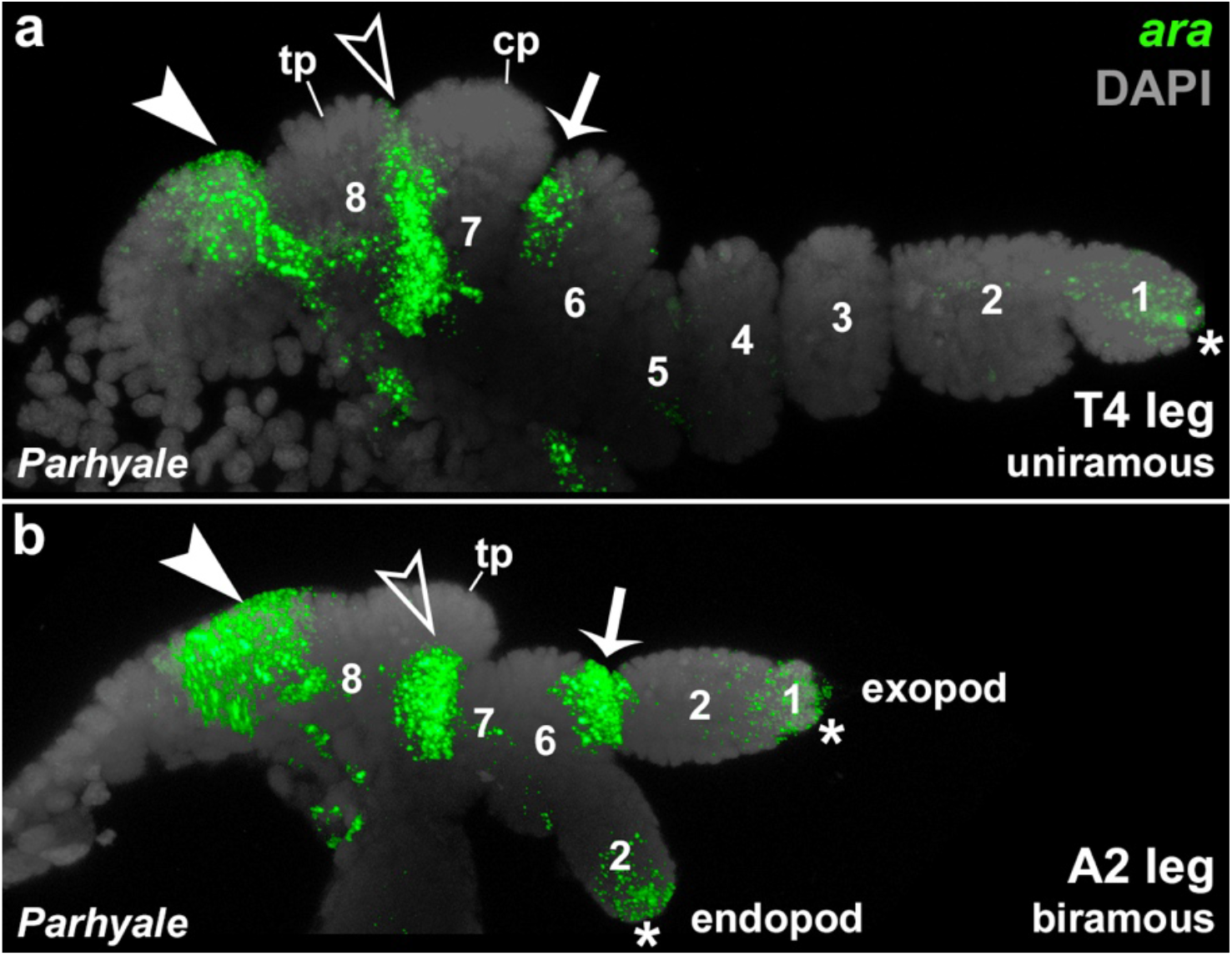
*ara* expression appears to mark the position of the exopod. In a uniramous *Parhyale* T4 leg that lacks an exopod, the leg segment 6 *ara* domain (arrow) forms a dot on the lateral side of the leg. However, in a biramous second abdominal (A2) *Parhyale* leg where an exopod emerges on the lateral side of leg segment 6, the *ara* dot is expanded to encompass the base of the exopod. *ara* armband on leg segment 8 (closed arrowhead). *ara* armband on lg segment 7 (open arrowhead). *ara* expression in distal tip of leg (^*^). The single segment on the *Parhyale* endopod could be leg segment 1 or 2 based on leg gene knockout and expression studies (results for abdominal legs are unpublished, but from same studies as in Bruce and Patel 2020^5^) but is here labeled as leg segment 2 for simplicity.

**Fig.S3.**
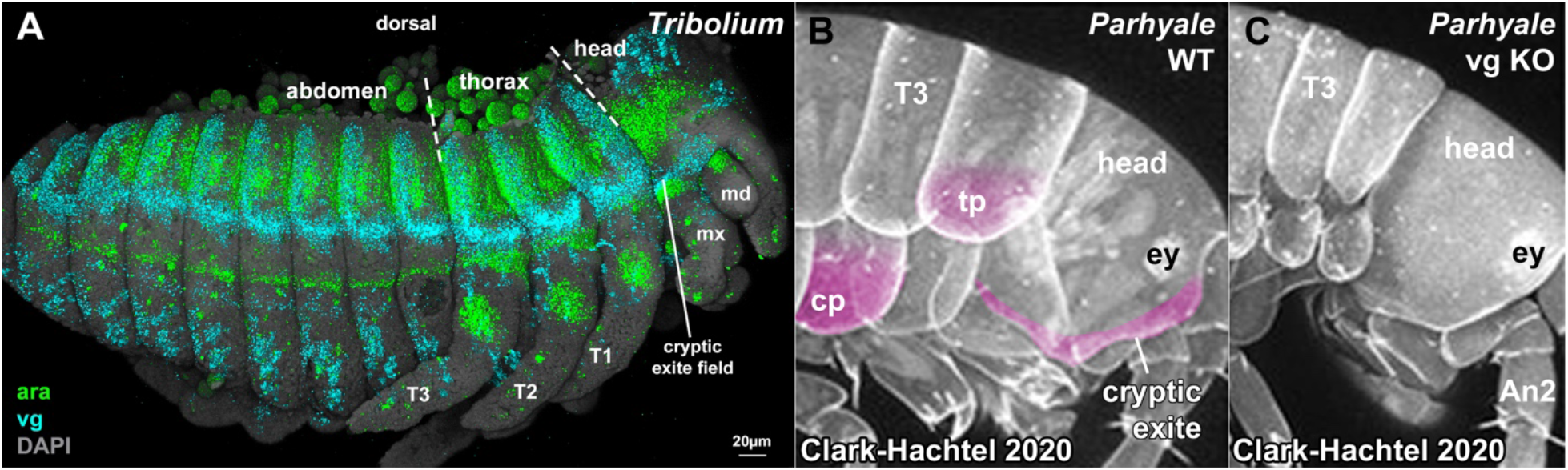
Cryptic exites in *Tribolium* and *Parhyale*. **A**. *Tribolium* embryo (lateral view). *vg* is expressed at the same register in all visible body segments. In all visible body segments, *vg* is bracketed dorsally and ventrally by *ara* expression domains. In thoracic legs T2 and T3, these *vg* expression domains pattern wings (the exites of the 8^th^ leg segment). In the abdominal segments, *vg* patterns the gin traps (larval defense structures), which are serially homologous to the wings^38^. In the head, as in the other body segments, *vg* is expressed in the same register and is bracketed by *ara*. Here, *vg* is expressed in the dorsal/proximal mandible (md) and maxilla (mx), and perhaps in the labium (homologous to crustacean mx2), which has migrated medial to the maxilla. **B**. WT Parhyale hatchling. A tergal plate (tp), coxal plate (cp), and the protrusion or flange on the head are shaded pink. **C**. When *vg* is knocked out, the tergal plate, coxal plate, and head flange are reduced (B, C, modified from Clark-Hachtel 2020^6^).

**Fig.S4.**
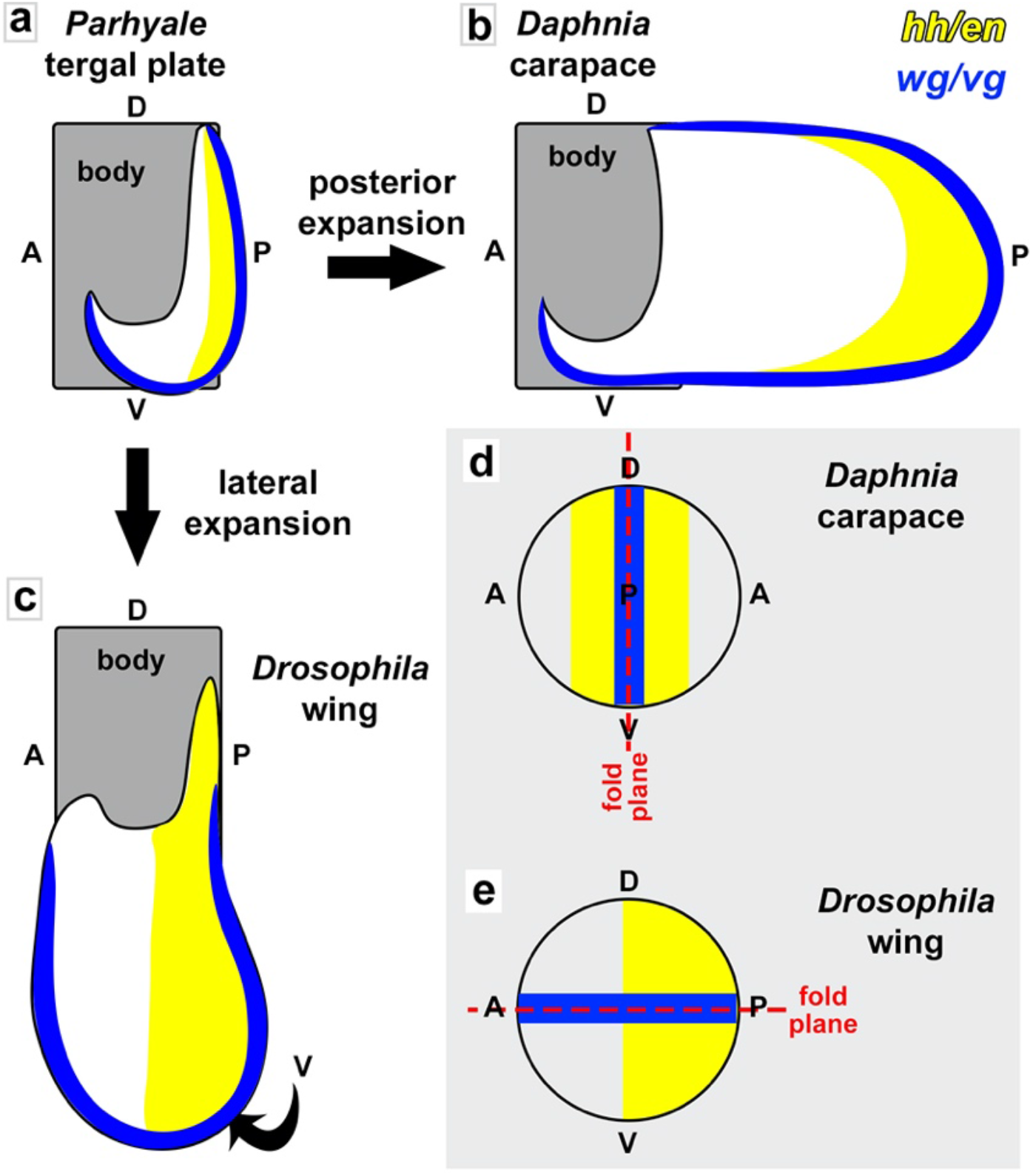
Summary of expression of *hh/en* and *wg*/*vg* in *Parhyale* tergal plate, *Daphnia* carapace, and *Drosophila* wing based on Shiga et al. 2017^2^ and Clark-Hatchel et al. 2020^6^. In *Parhyale* tergal plates (a), *vg* is expressed along the edge in a J-shape and *en* is expressed along the posterior edge^6^. If the posterior edge of a *Parhyale* tergal plate expanded posteriorly, then *en* and *wg* would be expressed in parallel in this expanded structure, as they are in the *Daphnia* carapace (b). If the ventral edge of a *Parhyale* tergal plate were to expand laterally, then *en* and *wg* would be expressed orthogonally to each other, as they are in insect wings (c). d – e reproduced from Shiga et al 2017 Fig 3M^2^.

## References

1. Müller, G.B., and Wagner, G.P. (1991). Novelty in evolution: restructuring the concept. Annu Rev Ecol Syst.

2. Shiga, Y., Kato, Y., Aragane-Nomura, Y., Haraguchi, T., Saridaki, T., Watanabe, H., Iguchi, T., Yamagata, H., and Averof, M. (2017). Repeated co-option of a conserved gene regulatory module underpins the evolution of the crustacean carapace, insect wings and other flat outgrowths. 120, 621–24.

3. Hu, Y., Schmitt-Engel, C., Schwirz, J., Stroehlein, N., Richter, T., Majumdar, U., and Bucher, G. (2018). A morphological novelty evolved by co-option of a reduced gene regulatory network and gene recruitment in a beetle. Proceedings of the Royal Society B: Biological Sciences 285, 20181373–9.

4. Fisher, C.R., Wegrzyn, J.L., and Jockusch, E.L. (2020). Co-option of wing-patterning genes underlies the evolution of the treehopper helmet. Nature Ecology & Evolution, 1–14.

5. Bruce, H.S., and Patel, N.H. (2020). Knockout of crustacean leg patterning genes suggests that insect wings and body walls evolved from ancient leg segments. Nature Ecology & Evolution 4, 1703–1712.

6. Clark-Hachtel, C.M., and Tomoyasu, Y. (2020). Two sets of candidate crustacean wing homologues and their implication for the origin of insect wings. Nat Ecol Evol.

7. Calman, W.T. (1909). A Treatise on Zoology. Part. VII. Appendiculata. Third Fascicle. Crustacea. (Lankester. London).

8. Fryer, G. (1996). The carapace of the branchiopod Crustacea. … Transactions of the Royal Society B: … 351:1703–1712.

9. Bruce, H.S. (2021). How to align arthropod leg segments. BioRxiv, 43.

10. Snodgrass, R.E. (1927). Morphology and mechanism of the insect thorax (City of Washington, Smithsonian institution).

11. Kobayashi, Y., Niikura, K., Oosawa, Y., and Takami, Y. (2013). Embryonic development of Carabus insulicola (Insecta, Coleoptera, Carabidae) with special reference to external morphology and tangible evidence for the subcoxal theory. J. Morphol. 274, 1323–1352.

12. Matsuda, R. (1970). Morphology and evolution of the insect thorax. Memoirs of the Entomological Society of Canada Volume 102, 5–431.

13. Tiegs, O.W. (1940). The embryology and affinities of the symphyla based on a study of Hanseniella agilis. Journal of Cell Science.

14. Boxshall, G.A., and Jaume, D. (2009). Exopodites, epipodites and gills in Crustaceans. Arthropod Systematics & Phylogeny, 1–27.

15. Ruiz-Losada, M., Blom-Dahl, D., Córdoba, S., and Estella, C. (2018). Specification and Patterning of Drosophila Appendages. JDB 6, 17–17.

16. Choi, H.M.T., Schwarzkopf, M., Fornace, M.E., Acharya, A., Artavanis, G., Stegmaier, J., Cunha, A., and Pierce, N.A. (2018). Third-generation in situhybridization chain reaction: multiplexed, quantitative, sensitive, versatile, robust. Development 145, dev165753–122.

17. Bruce, H.S., Jerz, G., Kelly, S., McCarthy, J., Pomerantz, A., Senevirathne, G., Sherrard, A. A, Sun, D., Wolff, C., and H Patel, N. (2021). Hybridization Chain Reaction (HCR) In Situ Protocol v1. protocols.io.

18. Ortega-Hernández, J., and Budd, G.E. (2016). The nature of non-appendicular anterior paired projections in Palaeozoic total-group Euarthropoda. Arthropod Structure and Development 45, 185–199.

19. Tomoyasu, Y. (2021). What crustaceans can tell us about the evolution of insect wings and other morphologically novel structures. Current Opinion in Genetics & Development 69, 48–55.

20. Medved, V., Marden, J.H., Fescemyer, H.W., Der, J.P., Liu, J., Mahfooz, N., and Popadić, A. (2015). Origin and diversification of wings: Insights from a neopteran insect. Proc. Natl. Acad. Sci. U.S.A. 112, 15946–15951.

21. Clark-Hachtel, C., Fernandez-Nicolas, A., Belles, X., and Tomoyasu, Y. (2021). Tergal and pleural wing-related tissues in the German cockroach and their implication to the evolutionary origin of insect wings. Evolution & Development.

22. Newman, W.A., and Knight, M.D. (1984). The Carapace and Crustacean Evolution—Arebuttal. Journal of Crustacean Biology 4, 682–687.

23. Boxshall, G.A. (2004). The evolution of arthropod limbs. Biol. Rev. 79, 253–300.

24. Davidson, E.H. (2006). Gene Regulatory Networks and the Evolution of Animal Body Plans. Science 311, 796–800.

25. Bowsher, J.H., and Nijhout, H.F. (2010). Partial co-option of the appendage patterning pathway in the development of abdominal appendages in the sepsid fly Themira biloba. Dev Genes Evol 219, 577–587.

26. Monteiro, A. (2012). Gene regulatory networks reused to build novel traits: Co-option of an eye-related gene regulatory network in eye-like organs and red wing patches on insect wings is suggested by optix expression. Bioessays 34, 181–186.

27. Lee, P.N., Callaerts, P., De Couet, H.G., and Martindale, M.Q. (2003). Cephalopod Hox genes and the origin of morphological novelties. Nature 424, 1061–1065.

28. Emlen, D.J., Corley Lavine, L., and Ewen-Campen, B. (2007). On the origin and evolutionary diversification of beetle horns. Proc. Natl. Acad. Sci. U.S.A. 104 Suppl 1, 8661–8668.

29. Moczek, A.P. (2009). On the Origins of Novelty and Diversity in Development and Evolution: A Case Study on Beetle Horns. Cold Spring Harbor Symposia on Quantitative Biology 74, 289–296.

30. Shubin, N., Tabin, C., and Carroll, S. (2009). Deep homology and the origins of evolutionary novelty. Nature 457, 818–823.

31. Shubin, N., Tabin, C., and Carroll, S. (1997). Fossils, genes and the evolution of animal limbs. Nature 388, 639–648.

32. Glassford, W.J., Johnson, W.C., Dall, N.R., Smith, S.J., Liu, Y., Boll, W., Noll, M., and Rebeiz, M. (2015). Co-option of an Ancestral Hox-Regulated Network Underlies a Recently Evolved Morphological Novelty. Developmental Cell 34, 520–531.

33. True, J.R., and Carroll, S.B. (2002). Gene co-option in physiological and morphological evolution. 32.

34. DiFrisco, J., Love, A.C., and Wagner, G.P. (2020). Character identity mechanisms: a conceptual model for comparative-mechanistic biology. Biol Philos 35, 44.

35. Stiassny, M. (2003). Atavism. In Keywords and concepts in evolutionary developmental biology Harvard University Press reference library., B. K. Hall and W. M. Olson, eds. (Harvard University Press).

36. Lewis, D.L., DeCamillis, M., and Bennett, R.L. (2000). Distinct roles of the homeotic genes Ubx and abd-A in beetle embryonic abdominal appendage development. Proc. Natl. Acad. Sci. U.S.A. 97, 4504–4509.

37. Clark-Hachtel, C.M., Linz, D.M., and Tomoyasu, Y. (2013). Insights into insect wing origin provided by functional analysis of vestigial in the red flour beetle, Tribolium castaneum. 110, 16951–16956.

38. Linz, D.M., and Tomoyasu, Y. (2018). Dual evolutionary origin of insect wings supported by an investigation of the abdominal wing serial homologs in Tribolium. 115, E658–E667.

39. Klüttgen, B., Dülmer, U., Engels, M., and Ratte, H.T. (1994). ADaM, an artificial freshwater for the culture of zooplankton. Water Research 28, 743–746.

40. Hansen, H.J. (1925). Studies on Arthropoda II. (Copenhagen: Gyldendalske Boghandel).

